# Molecular insights into nucleocapsid assembly and transport in Marburg and Ebola viruses

**DOI:** 10.1101/2025.05.13.653679

**Authors:** Yuki Takamatsu, Olga Dolnik, Ai Hirabayashi, Kenta Okamoto, Tomomi Kurashige, Hu Shangfan, Catarina Oda Harumi, Takeshi Noda

**Author notes:** Address correspondence to Yuki Takamatsu, Department of Virology, ITM-NU, Sakamoto1-12-4 Nagasaki, 852-8523, Japan.

## Abstract

Live-cell imaging enables visualization of the spatiotemporal dynamics of signals in cells. The intracytoplasmic movement of nucleocapsids is crucial during the lifecycle of enveloped viruses, but the molecular mechanisms governing their assembly and transport are not fully understood. Using a MARV live-cell imaging system, we identified three nucleocapsid proteins—NP, VP35, and VP24—that are necessary and sufficient to form transport-competent nucleocapsid-like structures (NCLSs). These findings are consistent with observations in Ebola virus (EBOV). Interestingly, despite incompatibility among these proteins, VP30 interacts with nucleocapsid proteins from both MARV and EBOV, supporting viral transcription and replication in heterologous systems. Furthermore, we show that the conserved PPxPxY motif at the C-terminus of NP regulates NP−VP30 interactions in both homologous and heterologous contexts, and is crucial for VP30 association with NCLSs. Because this motif is conserved across filoviruses, it represents a promising target for antiviral development. Our findings advance the understanding of nucleocapsid formation and offer new avenues for therapeutic intervention against MARV and EBOV.

## Introduction

The viral genome is encapsulated by proteins, forming nucleocapsids, to protect it from recognition by cellular defense mechanisms (Akira *et al*, 2006). In *Mononegavirales* virus infection, the newly synthesized nucleocapsid associated with viral RNA-dependent RNA polymerase complex is transported to the plasma membrane for virion formation and release (Schudt *et al*, 2013; Takamatsu *et al*, 2018a). Cryo-electron microscopy has revealed the high-resolution helical structure of RNA-bound nucleoprotein (NP) of Ebola virus (EBOV), Marburg virus (MARV), Cueva virus, Nipah virus, and measles virus (Desfosses *et al*, 2019; Fujita-Fujiharu *et al*, 2022; Hu *et al*, 2023; Ker *et al*, 2021; Sugita *et al*, 2018), which provides the molecular mechanisms driving and stabilizing the basic structure of assembled nucleocapsids. However, the complex structure of nucleocapsid, and the molecular mechanisms of NP-phosphoprotein (P) association, have been largely unrevealed. Measles virus P is mostly disordered, while three distinct interacting sites between it and NP contribute to efficient viral genome transcription (Guseva *et al*, 2019). In paramyxoviruses, polymerase and nucleoprotein are connected via the P protein, and a flexible conformational change may occur during RNA processing (Cox & Plemper, 2015; Guseva *et al*., 2019). The P protein is conserved in *mononegaviruses*, the roles of which are assumed to be divided on VP30 and VP35 in filoviruses (Biedenkopf *et al*, 2013; Biedenkopf *et al*, 2016; Kruse *et al*, 2018; Martinez *et al*, 2008; Modrof *et al*, 2002; Takamatsu *et al*, 2020b), whereas the interplay of nucleocapsid proteins during nucleocapsid assembly has been largely concealed.

MARV and EBOV belong to the family *Filoviridae* and have an approximately 19 kb non-segmented, single-stranded, negative-sense RNA genome. Although temporal epidemics have been reported over several years in central–western Africa, the first MARV epidemic was reported following an EBOV epidemic in 2021 in Guinea, Western Africa (WHO, 2021a, b), where the largest ever EBOV epidemic occurred during 2014–2016 with over 11,000 deaths (WHO, 2016). Since antiviral therapeutics for MARV and EBOV diseases are not well established, it is imperative to understand the molecular mechanisms of viral replication to establish countermeasures against these viruses. In this regard, revealing the intracellular dynamics of filoviruses is crucial, because nucleocapsid assembly and transport are prerequisites for virion formation.

Oligomerized NPs lead to the formation of inclusion bodies, which are sites for viral genome transcription, replication, and nucleocapsid synthesis (Huang *et al*, 2002; Kruse *et al*., 2018; Miyake *et al*, 2020; Noda *et al*, 2006; Watanabe *et al*, 2006). The core structure of filovirus nucleocapsids includes a nucleocapsid-like structure (NCLS) composed of NP, encapsulating single-stranded viral genomic RNA (Bharat *et al*, 2012; Muhlberger *et al*, 1998), together with the nucleocapsid protein VP24 and the polymerase cofactor VP35, both of which are essential structural elements that directly interact with NP to build a helical nucleocapsid approximately 800–1000 nm in length and 50 nm in diameter (Bharat *et al*., 2012; Huang *et al*., 2002; Takamatsu *et al*, 2020a; Watanabe *et al*., 2006). Viral polymerase L and transcription factor VP30 are also associated with the nucleocapsid (Biedenkopf *et al*., 2013; Hartlieb *et al*, 2003). An immunoelectron microscopy study of nucleocapsids demonstrated that NP, VP24, and VP35 are located from the center in this order, forming NCLS, and phosphorylation mediated VP30 association was observed at the peripheral region of NCLSs (Bharat *et al*, 2011; Takamatsu *et al*, 2022). Although the structural elements and viral protein properties are suggested to be highly conserved in filoviruses (Bharat *et al*., 2012; Wan *et al*, 2017), a detailed nucleocapsid proteins interaction has been yet to be clarified even in a recent cryo-electron microscopy analysis (Fujita-Fujiharu *et al*, 2025; Watanabe *et al*, 2024).

Here, we applied an EBOV live-cell imaging system to MARV and revealed essential viral factors, and compatibility of NC proteins during NC assembly and transport in MARV and EBOV. Moreover, common and distinct machinery of NC formation was identified between 2 viruses. Interestingly, PPxPxY motif mediates NP-VP30 interaction in filoviruses; however, in EBOV, it had no significant effect on transcription or replication. In contrast, VP30 lost its ability to support transcription and replication in MARV.

## Results

### Intracellular transport of MARV NCLSs nucleocapsids

The EBOV viral proteins VP30 and VP35 were utilized to visualize nucleocapsid and NCLS transport in live-cell imaging systems (Takamatsu *et al*., 2018a). GFP-conjugated VP30 and VP35 were analyzed and confirmed to be equivalent to their wild type in expression levels, subcellular localization, minigenomic functions and colocalization with other NCLS proteins (Fig. S1). Moreover, no differences in nucleocapsid transport characteristics were observed in MARV-VP30^GFP^- or MARV-VP35^GFP^-expressing cells (Fig. S2, Movies S1, S2) in accord with previous reports in EBOV infection (Dolnik *et al*, 2014; Schudt *et al*., 2013).

We previously reported that MARV NCLS transport can be visualized using a virus-like particle (VLP) system, expressing NP, VP35, VP24, L, VP40, GP, and minigenome, together with VP30^GFP^ (Takamatsu *et al*, 2019). Since VP40 and GP were not indispensable for EBOV NCLS transport (Takamatsu *et al*., 2018a), we transfected only MARV nucleocapsid components in Huh-7 cells and live-cell imaging was performed from 20 h post transfection (p.t.) (Fig. 1A). Smooth, long lines were observed in DMSO- or nocodazole-treated cells, whereas they disappeared in cytochalasin D-treated cells (Fig. 1B). The frequency of NCLSs translocated to a distance > 5 μm, as well as the mean velocity of NCLS transport, were drastically decreased in cytochalasin D-treated cells (Fig. 1C). In summary, NCLSs formed with NPs, VP35, VP30, VP24, L, and the minigenome exhibited long-distance transport via actin filaments, with characteristics similar to those of MARV nucleocapsid transport.

**Figure 1.**
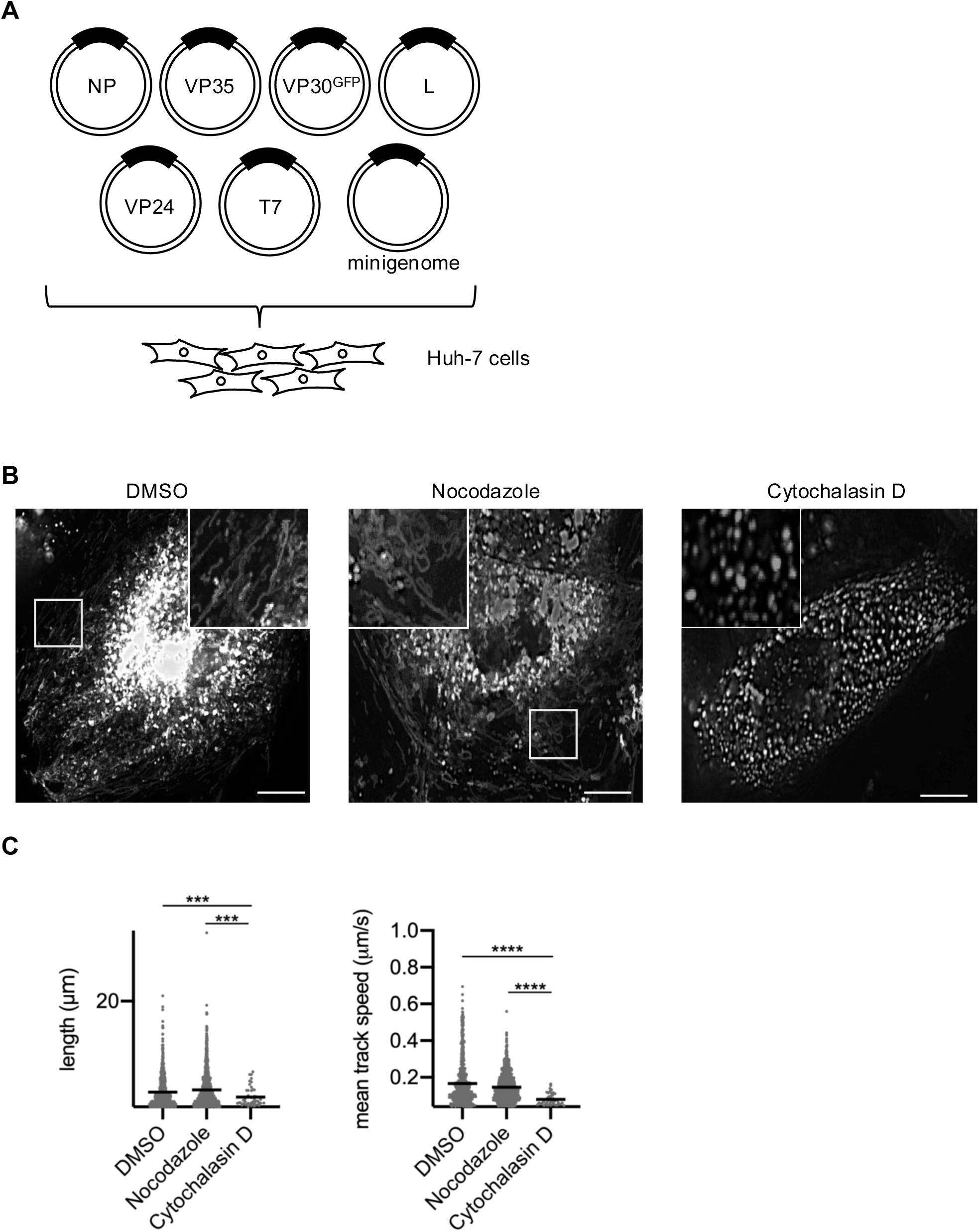
Live-cell imaging analysis of MARV nucleocapsid-like structure transport. (A) Schematic representation of the experimental settings for MARV NCLS transport. Huh-7 cells were transfected with plasmids encoding NP, L, VP35, VP24, MARV-specific minigenome, T7 polymerase, and VP30^GFP^. (B) Cells were observed from 20 h p.t. At 17 h p.t., different cytoskeleton-modulating drugs were added to the culture medium: 0.15% DMSO, 15 μM nocodazole, or 0.3 μM cytochalasin D, and the cells were incubated for an additional 3 h. The image shows the maximum-intensity projection of time-lapse images of cells, recorded for 3 min; images were captured every 3 s. The small boxed areas are enlarged at the four corners. Scale bars: 10 μm. (C) The detected signals were analyzed using the Imaris software. The length of the NCLS trajectories and velocity of NCLS transport were evaluated. The y-axis represents the signal counts. Lines indicate the mean ± SD. Asterisks indicate statistical significance; \*\*\**P* < 0.001 and \*\*\*\**P* < 0.0001.

### Identification of viral components necessary for MARV NCLS transport

Next, we applied a reductionist approach to identify viral factors required for NCLS transport. The experiments were repeated with the omission of each viral component (NP, VP35, VP24, L, or minigenome). Using live-cell imaging with MARV-VP30^GFP^ to track NCLS transport, we observed that even when polymerase L or the EBOV-specific minigenome was omitted, transport-competent NCLSs were still formed. In contrast, the omission of NP, VP35, or VP24 resulted in the failure to form transport-competent NCLSs (Fig. 2A). Next, the experiments were repeated using MARV-VP35^GFP^ to monitor the role of VP30 in NCLS transport. Although the presence of NP and VP24 is crucial for NCLS transport, VP30 is dispensable for NCLS transport (Fig. 2B). These data indicate that NP, VP35, and VP24 proteins are essential, whereas polymerase L, VP30, and the minigenome are not necessary for NCLS transport.

**Figure 2.**
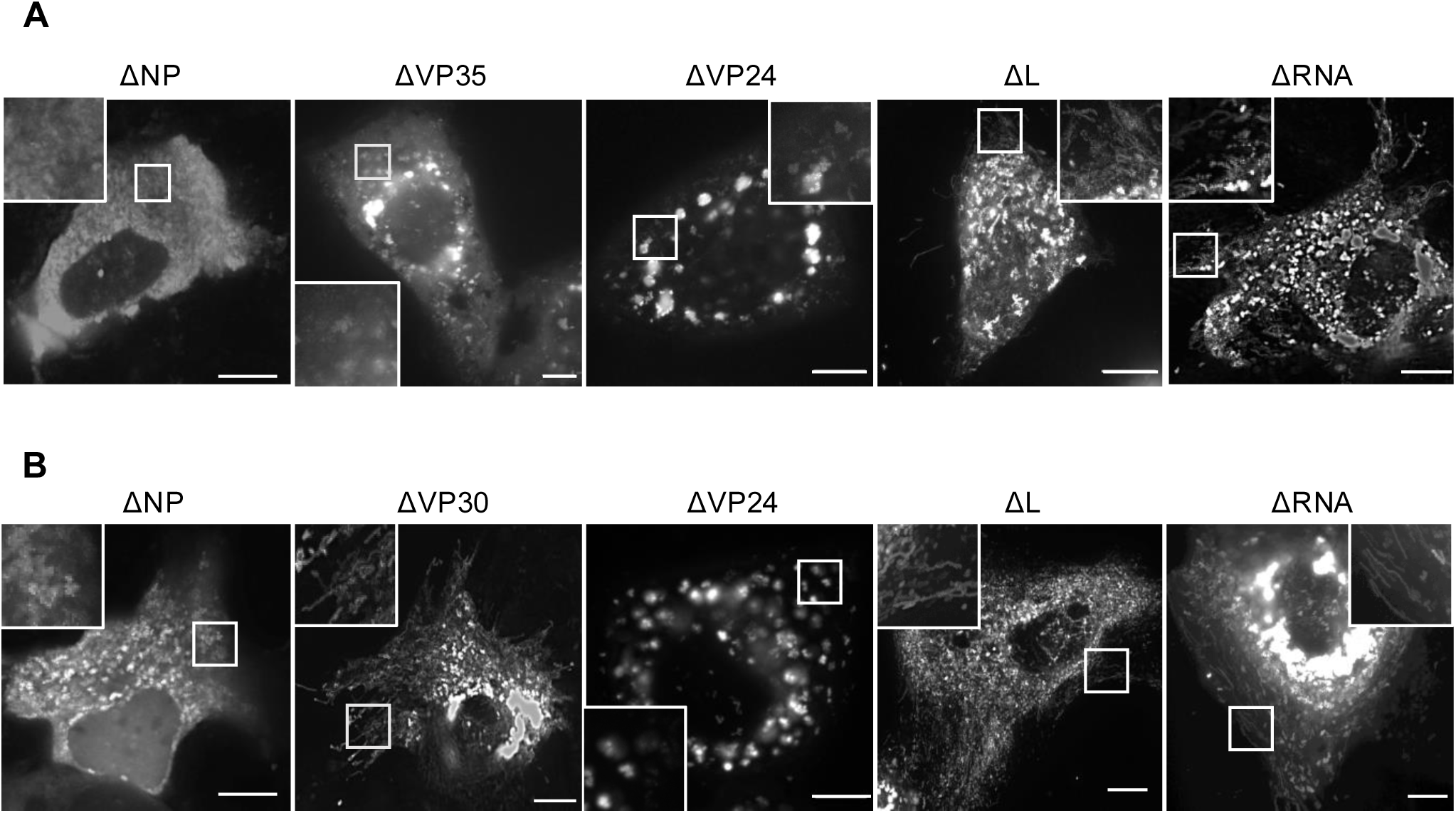
Determination of essential viral components for MARV NCLS transport. (A) Huh-7 cells were transfected with plasmids encoding VP30^GFP^, NP, VP35, VP24, L, MARV-specific minigenome, and T7 polymerase, except that one of the expression plasmids was omitted, as indicated. (B) Huh-7 cells were transfected with plasmids encoding VP35^GFP^, NP, VP30, VP35, VP24, L, MARV-specific minigenome, and T7 polymerase except that one of the expression plasmids was omitted, as indicated. Live-cell imaging analysis was started from 20 h p.t. The image shows the maximum-intensity projection of time-lapse images of cells recorded for 3 min; images were captured every 3 s. The small boxed areas are enlarged at the four corners. Scale bars: 10 μm.

### Transport characteristics of NP, VP35, and VP24 forming NCLSs

To confirm that these three components are sufficient to form transport-competent NCLSs, live cell imaging of NP, VP24 and VP35 together with VP35-GFP-expressing cells was started at 20h p.t. (Fig. 3A) Here again, we demonstrated that the frequency of NCLSs translocated > 5 μm, as well as the mean velocity of NCLS transport in nocodazole-treated cells (Fig. 3B-C, Movie S3), were drastically decreased in cytochalasin D-treated cells (Fig. 3B-C, Movie S4). The characteristics of NCLS transport were similar to those cells transfected with all nucleocapsid components (Fig. 1), and those of nucleocapsid transport in MARV-infected cells (Fig. S2). In summary, NP, VP35, and VP24 nucleocapsid proteins are necessary and sufficient to mediate NCLS transport.

**Figure 3.**
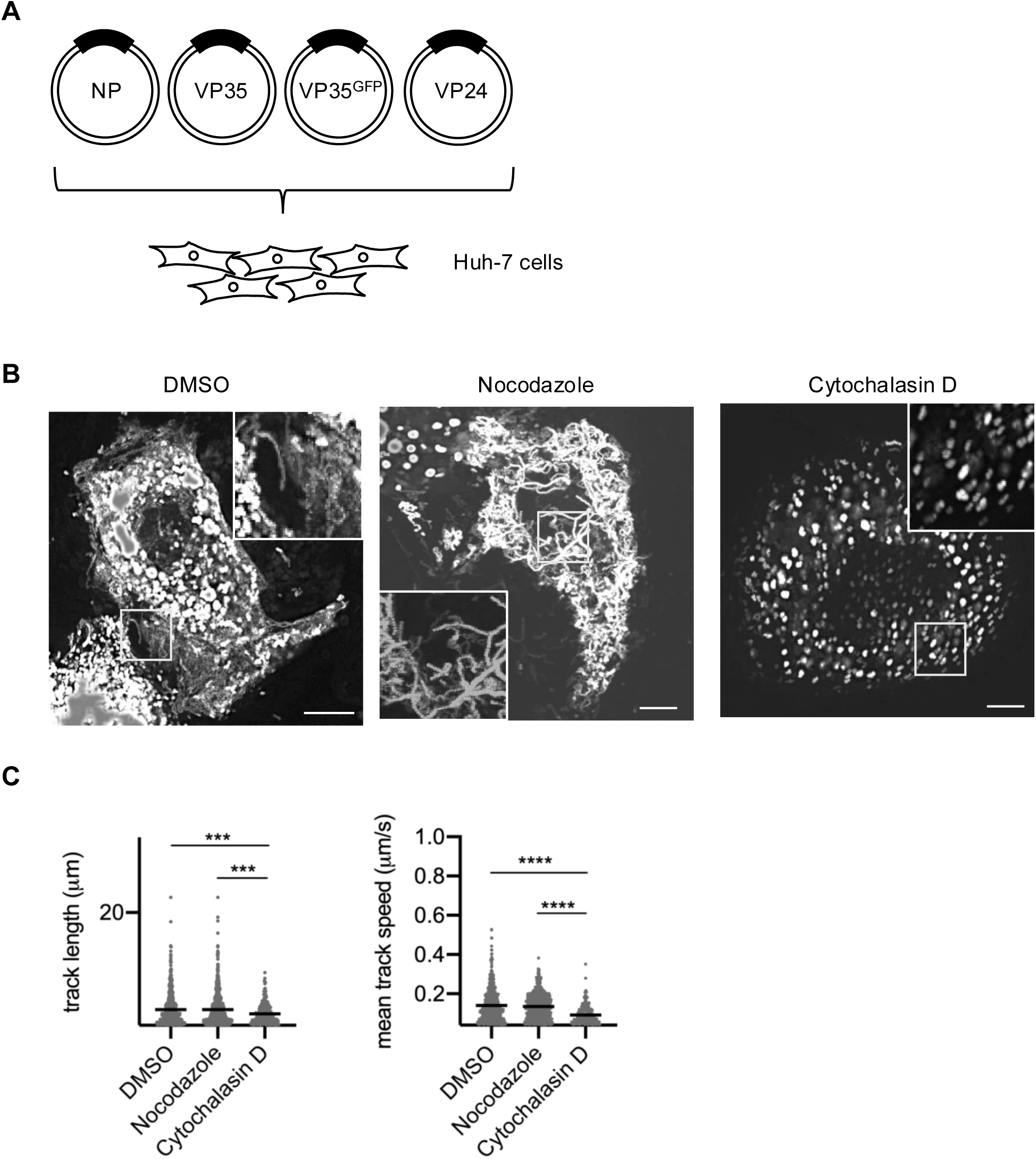
Live-cell imaging analysis of MARV NCLSs formed by NP, VP35 and VP24. (A) Schematic of experimental settings for NCLS transport, formed by NP, VP35 and VP24. (B) Huh-7 cells were transfected with the plasmids encoding NP, VP24, VP35^GFP^ and VP35. At 17 h p.t., different cytoskeleton-modulating drugs were added to the culture medium: 0.15% DMSO, 15 μM nocodazole, or 0.3 μM cytochalasin D for 3 h. Subsequently, cells were subjected to live-cell imaging analysis. The image shows the maximum-intensity projection of time-lapse images of cells, recorded for 3 min; images were captured every 3 s. The small boxed areas are enlarged at the four corners. Scale bars: 10 μm. (C) The detected signals were analyzed using Imaris software. The length of NCLS trajectories and velocity of NCLS transport were evaluated. The y-axis represents the number of signals. The lines indicate the mean ± SD. Asterisks indicate statistical significance; \*\*\**P* < 0.001 and \*\*\*\**P* < 0.0001.

### An exchange of NCLS proteins between MARV and EBOV

The helical structure of NCLS, which is formed by NP, VP35, and VP24, is well conserved in MARV and EBOV (Bharat *et al*., 2012; Wan *et al*., 2017), although interactions involving nucleocapsid-forming proteins between MARV and EBOV remain largely unexplored. To analyze the compatibility of nucleocapsid proteins between MARV and EBOV, we exchanged each protein in NP, VP35, VP24, and VP30 to observe the assembly and transport of NCLSs. In the MARV live-cell imaging system, EBOV NP, VP35, and VP24 were incompatible, whereas MARV-VP30 was exchangeable by EBOV-VP30 (Fig. 4A, Movie S5). EBOV-VP30 was also exchangeable by MARV-VP30 in the EBOV live-cell imaging system (Fig. 4B, Movie S6). These results indicate that VP30 proteins are structurally compatible with MARV and EBOV proteins.

**Figure 4.**
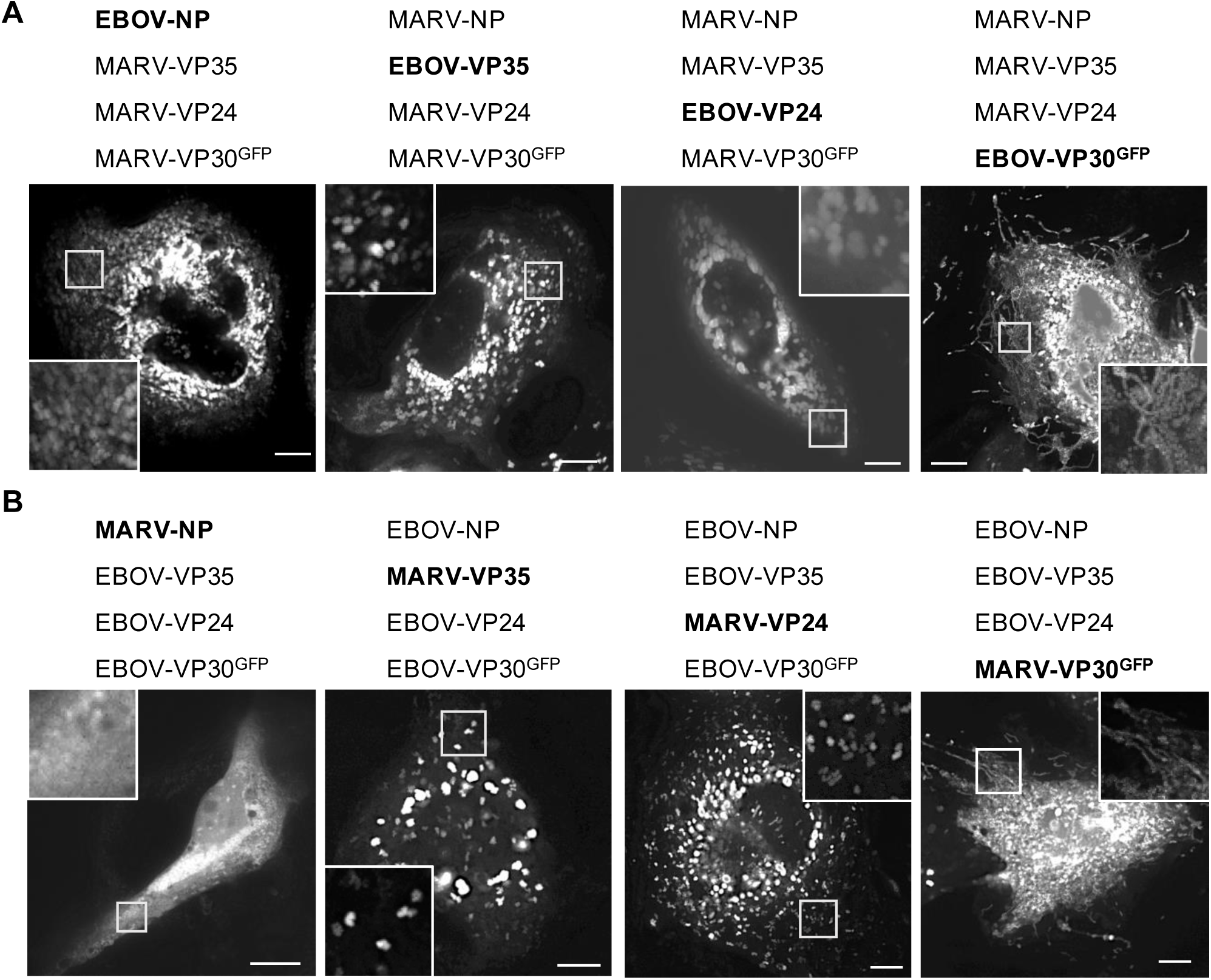
Live-cell imaging analysis of each NCLS protein replaced. (A) Huh-7 cells were transfected with plasmids encoding MARV proteins NP, VP35, VP24, and VP30^GFP^ but one protein was replaced with the corresponding EBOV protein (highlighted in bold). The indicated protein-coding plasmids were expressed. (B) Huh-7 cells were transfected with plasmids encoding EBOV proteins NP, VP35, VP24, and VP30^GFP^ but one protein was replaced with the corresponding MARV protein (highlighted in bold). The indicated protein-coding plasmids were expressed. Live-cell imaging analysis was started from 20 h p.t. The image shows the maximum-intensity projection of time-lapse images of cells, recorded for 3 min; images were captured every 3 s. The small boxed areas are enlarged at the four corners. Scale bars: 10 μm.

### Nucleocapsid protein replacement in minireplicon assays

Little information has been published regarding chimeric systems to evaluate genomic RNA replication among filoviruses (Muhlberger *et al*, 1999), and the magnitude and mechanisms of possible compatibility are unclear. Here, we replaced either NP, VP35, or VP30 in optimized minigenome transcription and replication assays (Hoenen *et al*, 2011; Wenigenrath *et al*, 2010). In the MARV minigenome assay, the exchange of MARV-NP with EBOV-NP and MARV-VP35 with EBOV-VP35 resulted in the abolishment of reporter activity close to the value of the negative control (in the absence of L protein). On the other hand, exchange of MARV-VP30 with EBOV-VP30 retained approximately 60% reporter activity, which was higher than that in the absence of MARV-VP30, which demonstrated around 40% reporter activity when the value of MARV-VP30 was set as 100% (Fig. 5A). In EBOV minigenome assays, the replacement of EBOV-VP30 with MARV-VP30 retained approximately 3-5% reporter activity, which was significantly higher than that in the absence of EBOV-VP30 with less than 1% reporter activity when the value of EBOV-VP30 was set as 100% (Fig. 5B). Interestingly, both MARV-VP30 and EBOV-VP30 were at least partially functional transcriptional activators in the heterologous replicon system.

**Figure 5.**
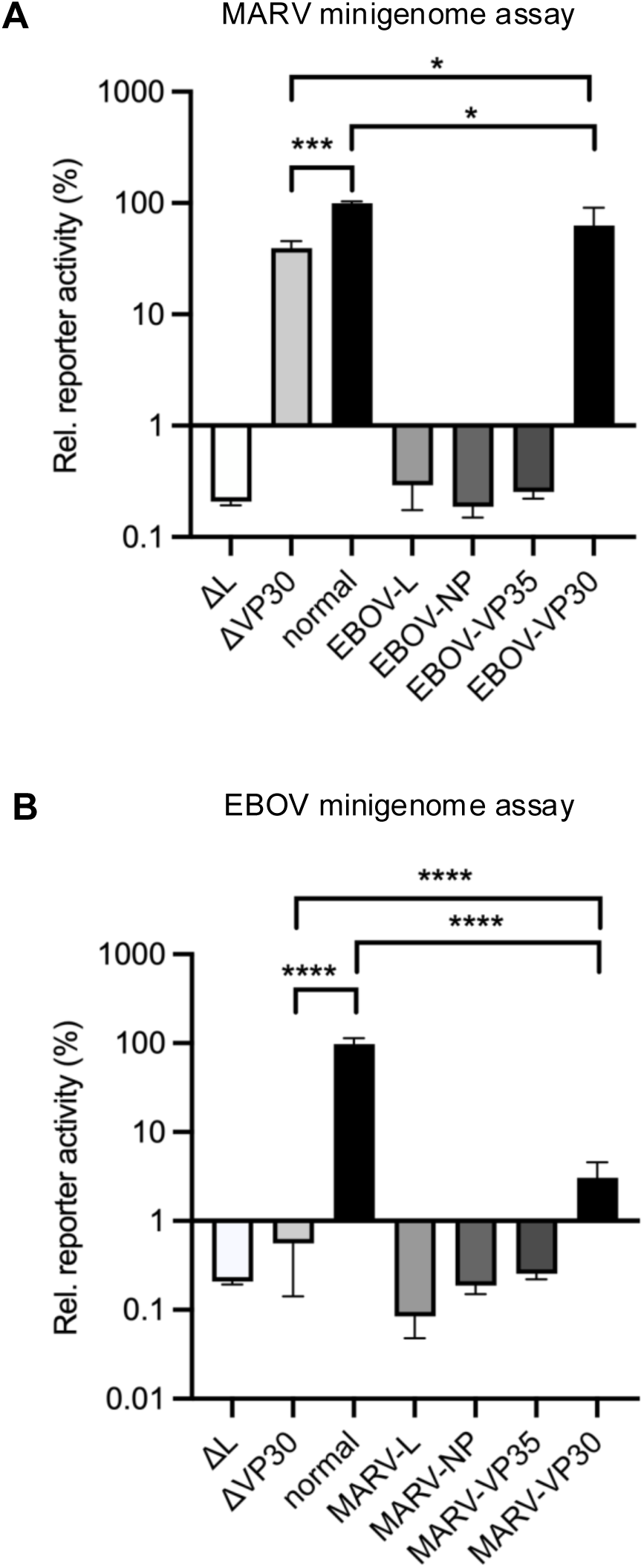
Minigenome assay with replacement of one viral protein. (A) HEK293 cells were transfected with MARV minigenome assay components where either NP, VP35, or VP30 was replaced with an EBOV protein. (B) HEK293 cells were transfected with EBOV minigenome assay components in which either NP, VP35, or VP30 was replaced with MARV protein. At 48 h p.t., cells were lysed, and reporter activity was measured. The value of normal MARV (A) or EBOV (B) minigenome component-transfected cells was set to 100%. The negative control (absence of L expression, ΔL) represented the background of the assay. Asterisks indicate statistical significance; \**P* < 0.05, \*\*\**P* < 0.001, \*\*\*\**P* < 0.0001.

### Interactions between NP and VP30

To reveal the molecular mechanisms underlying VP30-mediated transcriptional support activity and VP30-NCLS association in heterologous viruses, we performed immunofluorescence and immunoprecipitation assays. Both MARV-NP and EBOV-NP formed perinuclear inclusions, and both MARV-VP30 and EBOV-VP30 were diffusely distributed in the cytoplasmic regions when they were singly expressed (Fig. 6A-B, left lanes) (Becker *et al*, 1998; Modrof *et al*, 2001; Modrof *et al*., 2002). As previously described, MARV-VP30 accumulated in the MARV-NP-induced inclusions when co-expressed (Fig. 6A, middle lane) (Becker *et al*., 1998; Schudt *et al*., 2013). Similarly, EBOV-VP30 was localized in the EBOV-NP-induced inclusions when co-expressed (Fig. 6B, middle lane) (Biedenkopf *et al*., 2013; Modrof *et al*., 2001; Modrof *et al*., 2002). Notably, EBOV-VP30 also accumulated in MARV-NP-induced inclusions, and MARV-VP30 accumulated in EBOV-NP-induced inclusions (Fig. 6A-6B, right lanes).

**Figure 6.**
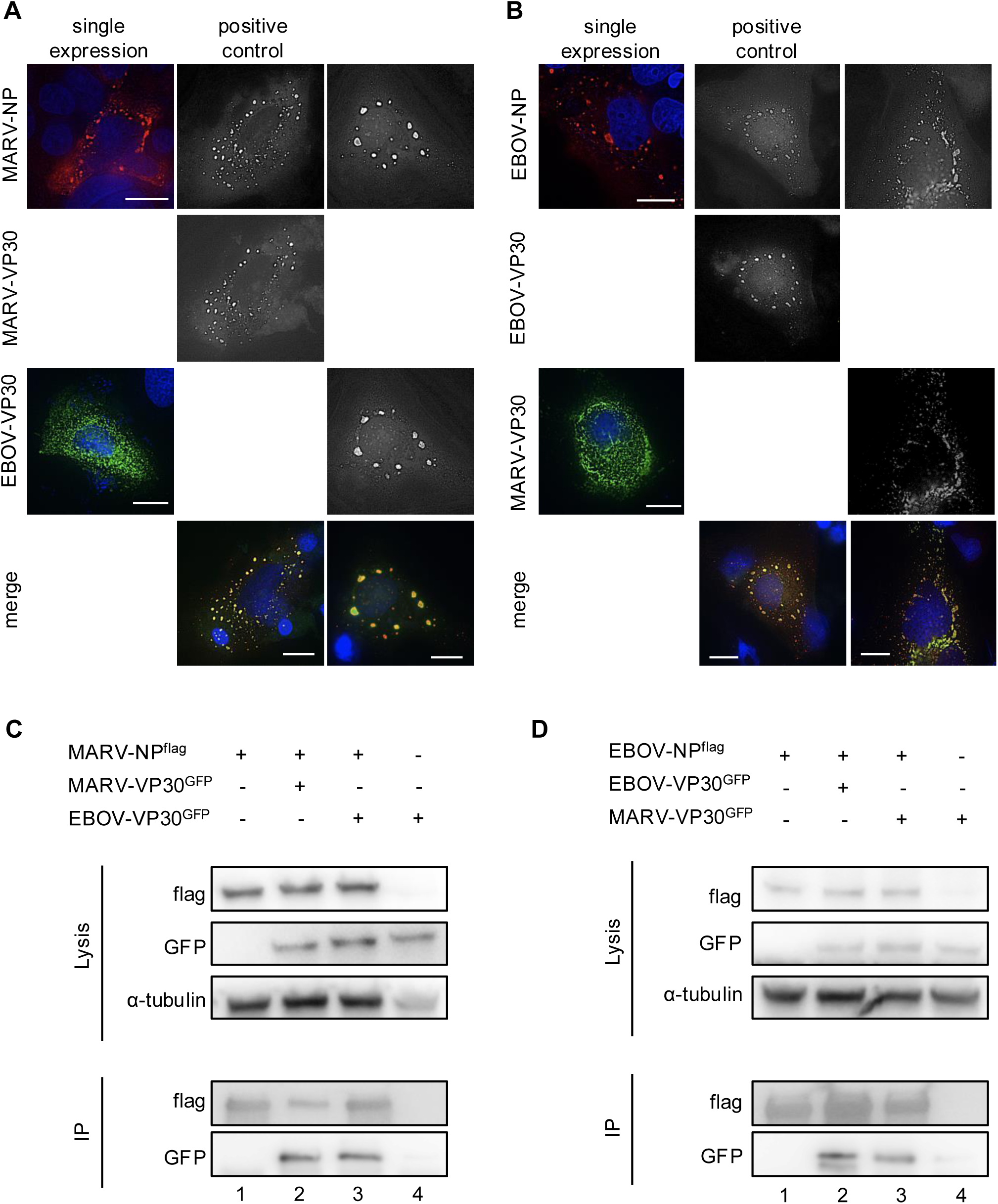
Homogenous and heterogenous NP−VP30 protein interactions. (A, B) Immunofluorescence analysis of Huh-7 cells. (A) Cells were transfected with a single plasmid: MARV-NP or EBOV-VP30 (left lane), a combination of plasmids MARV-NP and MARV VP30^GFP^ (middle lane), or MARV-NP and EBOV-VP30^GFP^ (right lane). (B) Cells were transfected with a single plasmid: EBOV-NP or MARV-VP30 (left lane), a combination of plasmids EBOV-NP and EBOV-VP30^GFP^ (middle lane), or EBOV-NP and MARV-VP30^GFP^ (right lane). Their intracellular distribution was analyzed using the corresponding NP-specific antibodies and/or autofluorescence. Scale bars: 10 μm. (C, D) Immunoprecipitation assays using MARV-NP^flag^ or EBOV-NP^flag^ in HEK293 cells. (C) MARV-NP^flag^ together with MARV-VP30^GFP^ or EBOV-VP30^GFP^-encoding plasmids were transfected into cells. (D) EBOV-NP^flag^ together with EBOV-VP30^GFP^ or MARV-VP30^GFP^-encoding plasmids were transfected into cells. At 48 h p.t., the cells were lysed and protein complexes were precipitated using mouse anti-Flag M2 agarose. An aliquot of the cell lysate (input) was collected before precipitation. Elution was achieved using SDS sample buffer. Western blot analysis was performed using Flag-, GFP-, and α-tubulin-specific antibodies. Lane numbers are indicated. Lane 4 is an absence of NP, representing non-specific interactions involving VP30^GFP^ and agarose beads.

Next, we performed immunoprecipitation assays with MARV-NP^flag^ and EBOV-NP^flag^. EBOV-VP30^GFP^ was precipitated using EBOV-NP^flag^, and MARV-VP30^GFP^ was precipitated using MARV-VP30^GFP^, as expected (Fig. 6C, lane 2, Fig. 6D, lane 2). Moreover, EBOV-VP30^GFP^ was precipitated by MARV-NP^flag^ and MARV-VP30^GFP^ was precipitated by EBOV-NP^flag^ (Fig. 6C-D, lanes 3). These results demonstrate that direct interactions between NP and VP30 bring about co-localization of these proteins in viral inclusions, suggesting that heterologous VP30 is partially functional in minigenome transcription and replication.

### PPxPxY motif-mediated NP-VP30 interactions

VP30 binds to the PPxPxY motif in NP, which regulates NP-mediated VP30 dephosphorylation (Kirchdoerfer *et al*, 2016; Kruse *et al*., 2018), which is a key modulator of transcriptional support activity in both EBOV and MARV (Biedenkopf *et al*., 2016; Modrof *et al*., 2002; Tigabu *et al*, 2018). To reveal interactions between VP30 and the NP PPxPxY motif, mutations were introduced into this motif (MARV-NP_ΔVP30_ and EBOV-NP_ΔVP30_, Fig. 7A). MARV-NP_ΔVP30_ exhibited reporter activity in the MARV minigenome assay, which was similar to that of the NP wild-type without VP30, regardless of the presence or absence of VP30 (Fig. 7B). In contrast, EBOV-NP_ΔVP30_ showed a decrease in reporter activity to the same level as that of the negative control when VP30 was not expressed; however, reporter activity was rescued by VP30 expression (Fig. 7C). As reported previously (Biedenkopf *et al*., 2013), VP30 in EBOV supports transcription and replication activity even when it loses its interaction with NP. Notably, VP30 does not exhibit transcription and replication support activity in MARV when it loses its interaction with NP.

**Figure 7.**
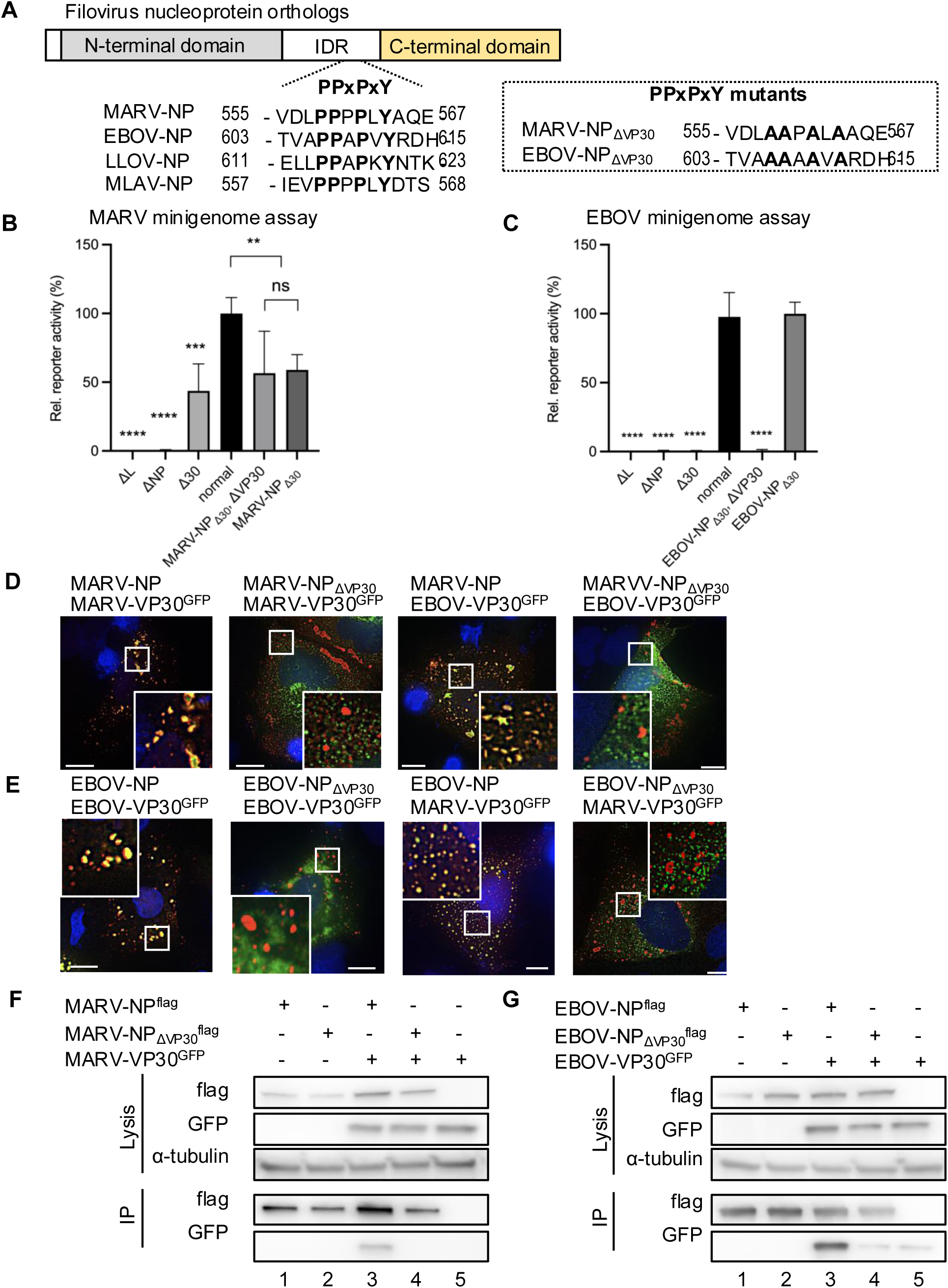
PPxPxY motif mediated interactions between NP and VP30 proteins. (A) Schematic representation of the regional orientation of the filoviral nucleoprotein orthologs. The N-terminal domain, intrinsically disordered region (IDR), and C-terminal domain are illustrated. Amino acid sequences of PPxPxY surrounding regions were selected and this motif is presented as bold and highlighted. The numbers indicate amino acid numbers counted from the N-terminal. PPxPxY motif mutants are enclosed in dotted boxes, named MARV-NP_ΔVP30_ and EBOV-NP_ΔVP30_. LLOV: *Lloviu cuevavirus*, MLAV: *Mengla dianlovirus*. (B, C) Minigenome assay in HEK293 cells. (B) In MARV minigenome assays, cells were transfected with minigenome components MARV-NP or MARV-NP_ΔVP30_ together with MARV-VP30 or EBOV-VP30. (C) In EBOV minigenome assays, cells were transfected with minigenome components EBOV-NP or EBOV-NP_ΔVP30_ together with EBOV-VP30 or MARV-VP30. At 48 h p.t., cells were lysed, and reporter activity was measured. The value of MARV-NP- or EBOV-NP-expressing cells was set to 100%. The mean and SD of three independent experiments are shown. Asterisks indicate statistical significance; **, *P* < 0.01; ***, *P* < 0.001; ****, *P* < 0.0001. (D, E) Immunofluorescence assay in Huh-7 cells. Cells were transfected with the indicated protein-encoding plasmids (D: MARV, E: EBOV). Intracellular distribution of proteins noted above the images was visualized using NP-specific antibodies and autofluorescence, and merged images were visualized. The small, boxed areas are enlarged at the four corners. Scale bars: 10 μm. (F, G) Immunoprecipitation assay in HEK293 cells. Cells were transfected with the indicated protein-encoding plasmids (F: MARV, G: EBOV). NP-encoding plasmids were fused with a FLAG tag, cells were lysed, and protein complexes were precipitated using mouse anti-FLAG M2 agarose at 48 h p.t. An aliquot of cell lysate (input) was collected before precipitation. Elution was achieved using SDS sample buffer. Western blot analysis was performed using FLAG-, GFP-, and alpha tubulin-specific antibodies.

Next, we employed confocal microscopy to visualize the formation of inclusion bodies and the localization of NP and VP30. This analysis demonstrated that MARV-NP_ΔVP30_ could form inclusion bodies but failed to incorporate either MARV-VP30 or EBOV-VP30 (Fig. 7D). Similarly, EBOV-NP_ΔVP30_ did not incorporate EBOV-VP30 or MARV-VP30 into the inclusions (Fig. 7E).

To clarify the interaction between NP and VP30 mediated by the PPxPxY motif, we conducted immunoprecipitation experiments. MARV-NP_ΔVP30_ lost its ability to interact with MARV-VP30 (Fig. 7F, lane 4), and EBOV-NP_ΔVP30_ was unable to precipitate EBOV-VP30 (Fig. 7G, lane 4).

These results indicate that the interactions between NP and VP30 are regulated by the conserved PPxPxY motif in NP, which is well conserved among filoviruses without affecting the formation of inclusions.

### PPxPxY motif and NCLS assembly, and VLP formation

Microscopic analyses were performed to investigate the role of the PPxPxY motif in NCLS assembly and transport. First, to analyze whether the NP mutants could form NCLSs, transmission electron microscopy was conducted. Tubular-like structures with electron-dense walls, representing condensed nucleocapsids, were detected in the presence of MARV-NP_ΔVP30_ (Fig. 8A, arrowheads) and EBOV-NP_ΔVP30_ (Fig. 8B, arrowheads).

**Figure 8.**
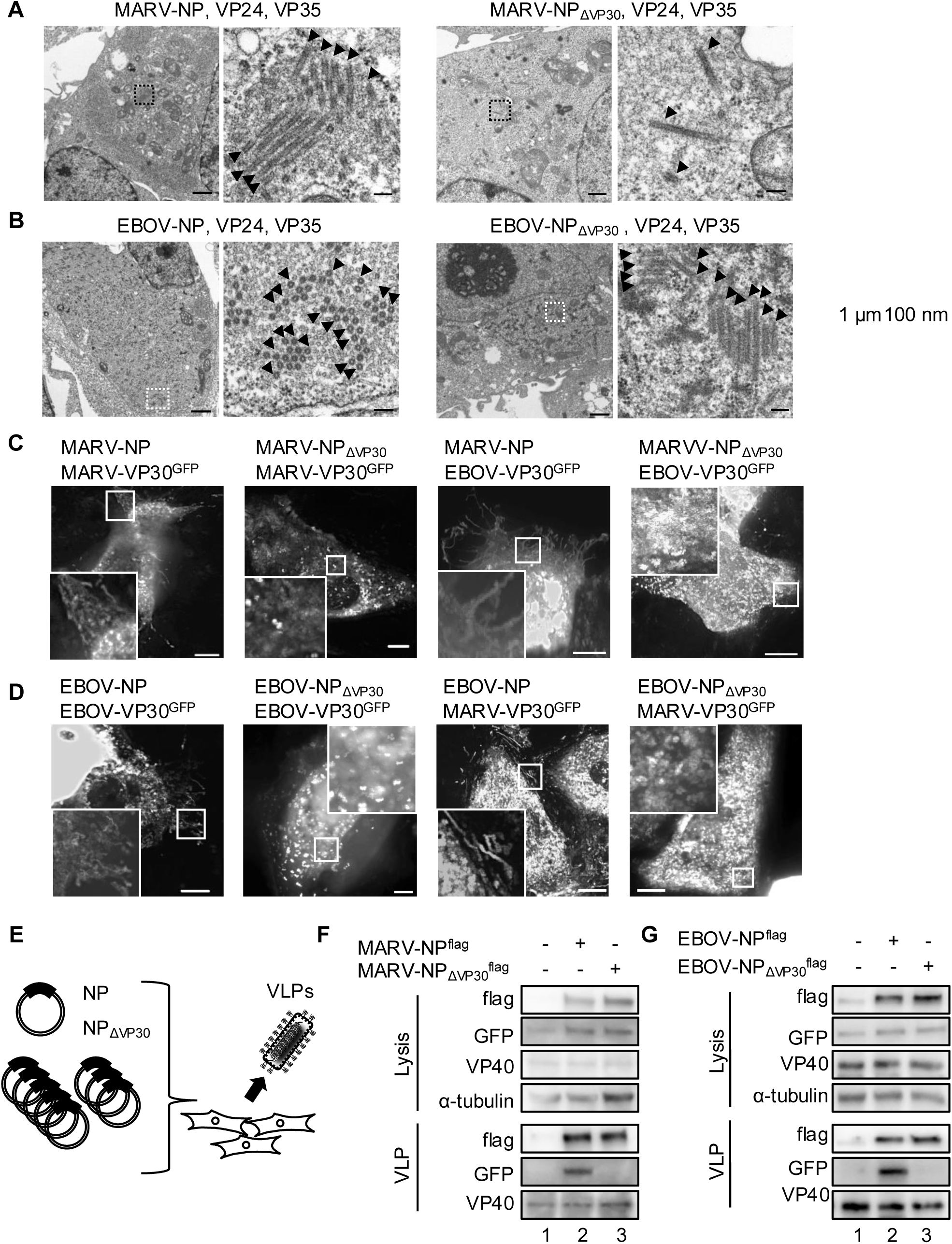
PPxPxY motif and VP30 proteins association to NCLSs, VLPs. (A, B) Huh-7cells expressing indicated proteins-coding plasmids, and were fixed at 48 h p.t. and processed for transmission electron microscopy analyses. The absence or presence of electron-dense walls in the tubular-like NCLSs is indicated by arrowheads on the transversal or longitudinal sections of NCLSs. The boxed areas in the left images are enlarged and are shown on the right. Scale bars are as follows: upper images 1 μm, lower images 100 nm. (C, D) Live-cell imaging in Huh-7 cells. Cells were transfected with the indicated proteins-encoding plasmids together with MARV-VP35 and MARV-VP24-encoding plasmids (C). Cells were transfected with the indicated proteins-encoding plasmids together with EBOV-VP35- and EBOV-VP24-encoding plasmids (D). Live-cell imaging analysis was started from 20 h p.t. The image shows the maximum-intensity projection of time-lapse images of cells, recorded for 2-3 min; images were captured every 2–3 s. The small boxed areas are enlarged at the four corners. Scale bars: 10 μm. (E) Schematic representation of MARV and EBOV virus-like particle (VLP) assay. (F, G) HEK293 cells were transfected with plasmids expressing all viral structural proteins and a MARV- or EBOV-specific minigenome encoding Renilla luciferase together with plasmid encoding firefly luciferase for normalization. The released VLPs were collected from supernatant, which was purified via sucrose cushion at 72 h p.t. Western blot was performed for viral proteins expression in producer cells and VLPs fractions (F: MARV, G: EBOV).

Next, we conducted live-cell imaging analyses of NCLS containing homologous or heterologous VP30-GFP in conjunction with NP or NP_ΔVP30_. As shown in Fig. 4, long sequential lines indicating transport-competent NCLSs were observed in MARV-NP- and EBOV-NP-containing NCLSs (Fig. 8C). In contrast, MARV-NP_ΔVP30_- and EBOV-NP_ΔVP30_-containing NCLSs demonstrated only diffusely distributed signals from both MARV and EBOV live-cell imaging systems (Fig. 8D).

Subsequently, we examined the process of virus-like particle (VLP) formation following NCLS transport. In the VLP assay, we transfected cells with NP, VP24, VP35, VP30, L, VP40, GP, a minigenome, and T7, and subsequently collected the VLPs secreted into the cell supernatant (Fig. 8E). Purified VLPs were assessed by western blotting. In both MARV and EBOV VLP assays, we observed the formation of VP40-containing VLPs, even in the absence of NP, whereas VP30 was undetectable (Fig. 8F-G, lanes 1). However, in the presence of wild-type NP, the incorporation of VP30 was confirmed; this was not observed in the NP mutants (MARV-NP_ΔVP30_ and EBOV-NP_ΔVP30_) (Fig. 8F-G, lanes 2 and 3).

In summary, mutations in the PPxPxY motif are crucial for VP30 association with NCLSs and for VP30 incorporation into VLPs in both MARV and EBOV.

### PPxPxY motif and NC protein interactions

The association between NP and VP35 was reported upon NP binding peptide (20-48 a.a. of VP35) to the N-terminal side of NP (residues 25-457), and the basic patch of VP35 (residues 220-251) to the central domain of NP (Kirchdoerfer *et al*., 2016; Leung *et al*, 2015; Leung *et al*, 2010; Miyake *et al*., 2020; Zhu *et al*, 2017). Regarding the NP-VP24 interaction, two molecules of VP24 bind to two molecules of NP in distinct configurations, and mutations in NP_R132A_ and NP_H196A_ inhibit the formation of NCLS (Fujita-Fujiharu *et al*., 2025). Given that the structure of the C-terminal region of NP and the interactions among NCLS proteins are largely unknown, we used AlphaFold2 and AlphaFold3 (Abramson *et al*, 2024; Jumper *et al*, 2021) to predict the binding of PPxPxY peptides to VP35 and VP24. Confident structural predictions were obtained only for MARV-VP35, EBOV-VP35, and EBOV-VP24 (pLDDT > 80, Fig. S3A-C). All the predicted structures indicated that both MARV and EBOV VP35 interacted with the PPxPxY peptide in a similar structural region. These results suggest that the PPxPxY peptide may interact with the positively charged VP35 surface in both MARV and EBOV (Fig. S3D-E). However, a protein complex was not predicted for the peptide with an alanine substitution. Additionally, we were unable to predict a highly confident binding of the PPxPxY peptide to MARV-VP24; in contrast, both PPxPxY and the alanine substitution mutant peptides appeared to interact with EBOV-VP24 (Fig. S3C).

To reveal the NP-PPxPxY motif and VP35 interaction, microscopy and immunoprecipitation assays were used. Immunofluorescence microscopy analyses revealed that MARV-NP_ΔVP30_ recruited MARV-VP35 to these inclusions (Fig.S4A). Similarly, EBOV-NP_ΔVP30_ recruits EBOV-VP35 (Fig. S4B). Next, we conducted live-cell imaging analyses of NCLSs containing either NP or NP_ΔVP30_ along with their respective VP35-GFP. Both MARV-NP_ΔVP30_ and EBOV-NP_ΔVP30_ exhibited transport-competent NCLSs (Fig. S4C-D). Finally, we performed immunoprecipitation assays, demonstrating that both MARV-NP_ΔVP30_ and EBOV-NP_ΔVP30_ successfully precipitated their respective VP35 (Fig. S4E-F, lanes 4). These results suggest that the PPxPxY motif does not govern the interactions among NCLS-forming proteins; rather, it regulates the interaction between NP and VP30 in a localized manner.

In summary, VP30 exhibits compatibility between MARV and EBOV, with transcription and replication activities partially sustained by heterologous VP30 (Fig. 9A). The binding of NP to VP30 is regulated by the PPxPxY motif; when a mutation is introduced, VP30 is unable to bind to NP. Consequently, there was no association between VP30 and NCLS (Figure 9B). Conversely, when the motif is intact, heterologous VP30 can bind to NCLS, facilitating intracellular transport (Fig. 9C).

**Figure 9.**
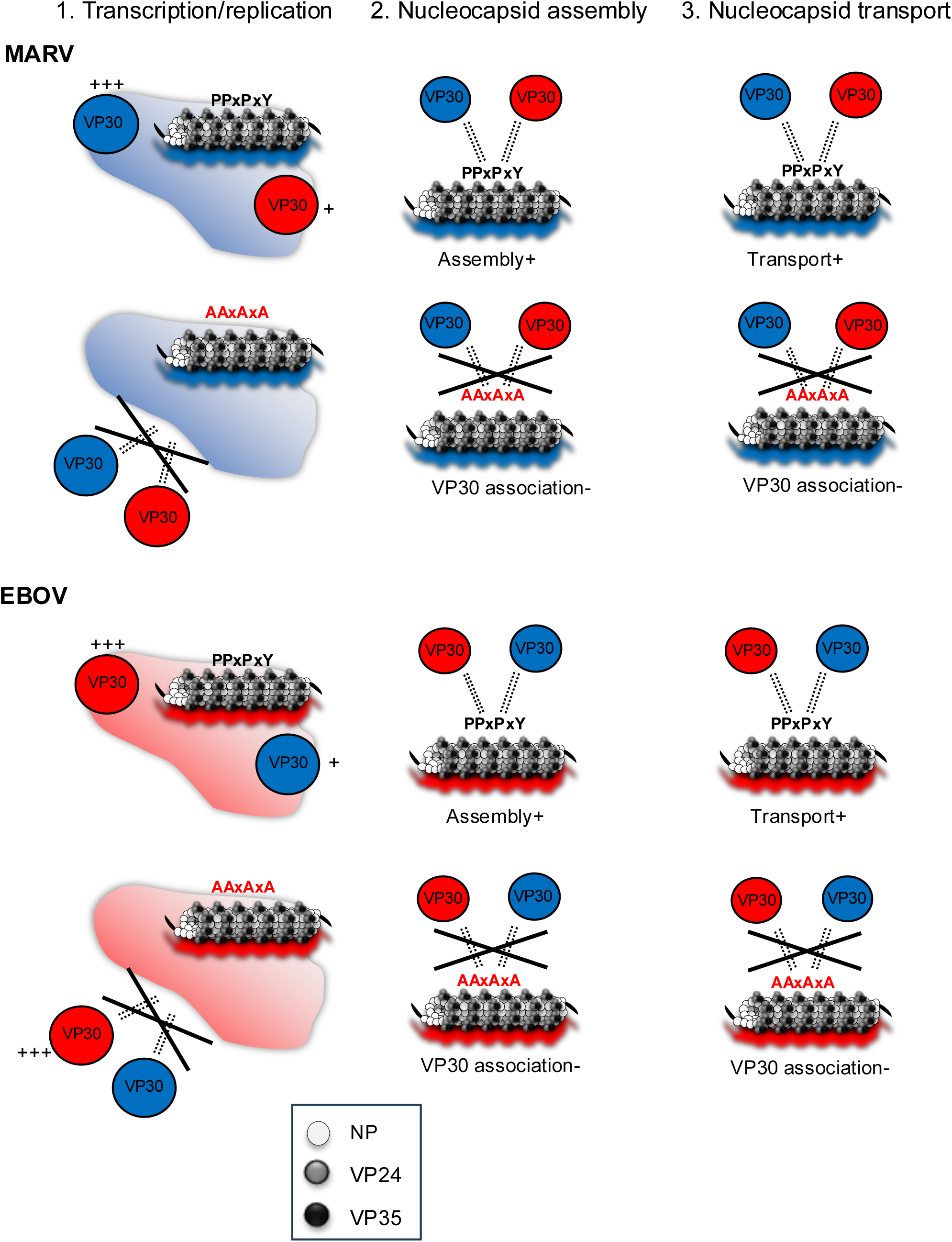
Role of the PPxPxY motif of NP during NCLS assembly in MARV and EBOV. This figure illustrates the current understanding of NCLS assembly and VP30-NCLS associations across three key processes: (1) viral genome transcription/replication, (2) nucleocapsid assembly, and (3) nucleocapsid transport. (1) In filoviruses, NP forms perinuclear inclusions where viral genome transcription/replication and nucleocapsid synthesis occur. VP30 co-localizes with heterologous NP-forming inclusions and partially restores heterologous minigenome transcription/replication. (2) Mutations in the PPxPxY motif regulate NP-VP30 interactions in both MARV and EBOV. (3) VP30 is associated with heterologous NP, facilitating its linkage to NCLS. This indicates that NCLS is initially formed by NP, VP35, and VP24, with VP30 association occurring during or after NCLS assembly.

## Discussion

In the present study, we revealed that the core structure of nucleocapsids, NCLSs, is formed from NP, VP35, and VP24 (Fig. 1-3) in agreement with EBOV, using live-cell imaging systems based on the ectopic expression of fluorescently labelled viral proteins in MARV. Based on these observations, we characterized the nucleocapsid compatibility between MARV and EBOV and sought to reveal the molecular interface between the nucleocapsid-forming proteins.

The characteristic filoviral NP-RNA helical complex provides a scaffold for nucleocapsid formation, which is responsible for the transcription and replication of viral RNAs (Bharat *et al*., 2012; Fujita-Fujiharu *et al*., 2022; Hu *et al*., 2023; Sugita *et al*., 2018). Previously MARV-VP30 was reported to be partially functional in the transcription and replication of EBOV chloramphenicol acetyltransferase reporter assays (Muhlberger *et al*., 1999). Our results demonstrated that EBOV-VP30 also supported transcription and replication of MARV minigenome (Fig. 5B), indicating that filoviruses have conserved machinery for VP30-nuclecoapsid associations. On the other hand, VP30 plays different roles in recombinant viruses production; MARV was rescued by EBOV VP30 instead of MARV VP30 expressions, whereas EBOV was rescued only by EBOV VP30 (Enterlein *et al*, 2006). VP30 is a phosphoprotein, and its dephosphorylation is crucial for transcriptional support in EBOV rather than in MARV (Biedenkopf *et al*., 2013; Kruse *et al*., 2018; Tigabu *et al*., 2018). The LxxIxE and PPxPxY motifs located adjacent to the C-terminus of NP are conserved in filoviruses (Kruse *et al*., 2018). In EBOV, VP30 binds to NP at the PPxPxY motif, and VP30 is dephosphorylated by PP2A, which is recruited by the LxxIxE motif (Kruse *et al*., 2018). Notably, mutations involving the PPxPxY motif did not significantly affect the transcriptional support activity in either the MARV or EBOV minigenome assays (Fig. 7B, 7C). Defective interactions involving NP-VP30 proteins do not cause defective viral genome transcription/replication in EBOV (Biedenkopf *et al*., 2013; Kirchdoerfer *et al*., 2016), indicating that a high-affinity interaction between NP and VP30 is not strictly required for viral RNA synthesis, and minimal binding of these proteins mediates viral RNA synthesis in EBOV (Xu *et al*, 2017). In contrast, the transcription and replication activity of MARV slightly decreased due to mutation of the PPxPxY motif. However, this reduction was consistent with the levels observed in the absence of VP30 in wild-type NP, suggesting that this phenomenon reflects the influence of VP30 rather than the activity of the motif itself in transcription and replication.

In filoviruses, nucleocapsids form a left-handled helix with an inner nucleoprotein layer decorated with protruding arms composed of VP24 and VP35 (Bharat *et al*., 2012; Noda *et al*., 2006; Wan *et al*., 2017). Even in a recent model, the intrinsically disordered C-terminal region (IDR) of NP (aa 450-600), which is critical for nucleocapsid formation in the presence of VP24 and VP35, has been unclear due to its flexibility and insufficient resolution of the EM map (Fujita-Fujiharu *et al*., 2025; Watanabe *et al*., 2024). This study reveals a common aspect of filoviral nucleocapsid assembly, that is, heterologous VP30 associates with NP and supports transcription/replication in the inclusions in both MARV and EBOV (Fig. 5, 6). Intrudingly, the PPxPxY motif regulates the interaction with NP-VP30, but does not affect the assembly and transport of NCLSs (Fig.8A-B, Fig.S4C-D), Consequently, introducing mutations in this motif inhibits the association of VP30 to VLPs (Fig.8F, 8G). Noteworthy, it has been reported that transcription and replication are inhibited by the addition of competitive peptides against this motif in EBOV minigenome (Xu *et al*., 2017). Moreover, PPxPxY motif-bearing proteins, such as RBBP6, hnRNPUL1, and PEG10 modulate EBOV transcription and replication through discrete mechanisms (Batra *et al*, 2021), highlighting the importance of this motif in EBOV replication. Given that research on this motif in MARV has not yet been reported, it remains a topic for future investigation.

Using AlphaFold2/3 prediction (Abramson *et al*., 2024; Jumper *et al*., 2021), the PPxPxY peptide may bind to the positively charged surface of VP35, both in MARV and EBOV, although no such protein complexes are predicted when peptides hold alanine substitutions (Fig. S3). In EBOV, amino acid residues R225, H240, K248, and K251 of VP35 are reported to be important for its NP interaction (Prins *et al*, 2010), which forms the first basic patch (amino acid residues 222-251) (Miyake *et al*., 2020). Interestingly, residues Q241, Q244, and K248 located in this basic patch have been shown to be part of the PPxPxY-binding motif of the NP, but mutations in this motif did not affect the NP-VP35 interaction or the formation of NCLS (Fig. 8A-B, S4).

In conclusion, our study demonstrated that the PPxPxY motif not only regulates the binding between NP and VP30 but also influences the association of VP30 with NCLS. Interestingly, the interaction between NP and VP30 through the PPxPxY motif is regulated in a somewhat permissive manner, exhibiting compatibility between MARV and EBOV. This suggests a potential for developing drugs that inhibit the replication of a wide range of filoviruses by targeting non-specific binding through this motif.

## Materials and methods

### Cells and Viruses

Huh-7 and HEK293 cells were maintained at 37°C and 5% CO_2_ in Dulbecco’s modified Eagle’s medium (DMEM; Life Technologies) supplemented with 10% (v/v) fetal bovine serum (FBS; PAN Biotech), 5 mM L-glutamine (Q; Life Technologies), 50 U/mL penicillin, and 50 μg/mL streptomycin (PS; Life Technologies). MARV (Musoke accession no. DQ217792.1, GenBank) and recombinant MARV were propagated on VeroE6 cells as described previously (Krahling *et al*, 2010). All work with infectious viruses was performed in a BSL-4 facility at Philipps-Universität Marburg following national legislation and guidelines.

### Plasmids and transfection

Plasmids encoding the MARV proteins pCAGGS-MARV-NP, pCAGGS-MARV-VP35, pCAGGS-MARV-VP30, pCAGGS-MARV-VP24, pCAGGS-MARV-L, pCAGGS-MARV-VP40, and pCAGGS-MARV-GP, as well as the T7-driven MARV minigenome encoding *Renilla* luciferase and pCAGGS-T7 polymerase, were used as previously described (Hoenen *et al*., 2011;Wenigenrath *et al*., 2010). The plasmids encoding MARV-VP30^GFP^ and MARV-VP35^GFP^ fusion proteins have been described previously (Schudt *et al*., 2013; Takamatsu *et al*, 2018b). Plasmids encoding EBOV proteins (pCAGGS-EBOV-NP, pCAGGS-EBOV-VP35, pCAGGS-EBOV-VP30, pCAGGS-EBOV-VP24, and pCAGGS-EBOV-L) and the T7-driven EBOV minigenome encoding *Renilla* luciferase were prepared as described previously (Biedenkopf *et al*., 2013; Hoenen *et al*, 2006).

The cloning procedure for plasmids with mutations introduced at the PxPPxY motif with Flag-tagged plasmids (pCAGGS-MARV-NP^flag^, pCAGGS-MARV-NP_ΔVP30_^flag^, pCAGGS-EBOV-NP^flag^, and pCAGGS-EBOV-NP_ΔVP30_^flag^) has been described previously (Takamatsu *et al*., 2020a).

Transfection was performed in Opti-MEM without phenol red (Life Technologies) using TranSIT (Mirus) according to the manufacturer’s instructions.

### SDS-PAGE and western blot analysis

SDS-PAGE and western blot analyses were performed as previously described (Biedenkopf *et al*., 2013; Kolesnikova *et al*, 2004). Protein detection was performed using Image Lab™ software (Bio-RaD) or Image Reader LAS-3000 (Fujifilm) for HRP-conjugated secondary antibodies, as indicated in the antibodies section below.

### Immunofluorescence analysis and confocal laser scanning microscopy

Immunofluorescence analyses were performed as previously described (Dolnik *et al*, 2010; Kolesnikova *et al*, 2012). Microscopic images were acquired using a Leica SP5 confocal laser scanning microscope with a 63× oil objective, Olympus FV3000 microscope with a 100× oil objective, or Keyence BZ-X810 microscope with a 100× oil objective (Olympus). Cells were grown on µ-Slide 8 or 12 wells (ibidi) and fixed with 4% paraformaldehyde 20 h post-transfection. Nuclear staining was performed using Hoechst33342 (Dojindo).

### Antibodies

The following primary antibodies were used for immunofluorescence analysis: mouse anti-MARV-NP (Wenigenrath *et al*., 2010), rabbit anti-MARV-NP (IBT), rabbit anti-MARV-VP30 (Wenigenrath *et al*., 2010), guinea pig anti-MARV-VP35 (Wenigenrath *et al*., 2010), rabbit anti-MARV-VP24 (Wenigenrath *et al*., 2010), chicken anti-EBOV-NP (Biedenkopf *et al*., 2016), rabbit anti-EBOV-NP (IBT), and rabbit anti-EBOV-VP30 (Biedenkopf *et al*., 2013). The corresponding secondary antibodies were donkey anti-mouse Alexa488 (abcam), donkey anti-mouse Alexa594 (abcam), donkey anti-mouse-IRDye 680RD (LI-COR), donkey anti-mouse Alexa488 (abcam), donkey anti-rabbit Alexa594 (abcam), donkey anti-rabbit IRDye 680RD (LI-COR), goat anti-guinea pig Alexa594 (abcam), goat anti-chicken Alexa594 (Thermo Fisher Scientific), and donkey anti-chicken IRDye 680RD (Li-COR).

The following primary antibodies were used for western blot analysis: mouse anti-MARV-NP monoclonal antibody (Dolnik *et al*., 2014), rabbit anti-MARV-NP antibody (IBT), mouse anti-MARV-VP40 monoclonal antibody (Dolnik *et al*., 2014), chicken anti-EBOV-NP polyclonal antibody (see above), rabbit anti-EBOV antibody (IBT), rabbit anti-EBOV-VP40 antibody (IBT), rabbit anti-GFP antibody (Rockland), mouse anti-FLAG M2 monoclonal antibody (Sigma-Aldrich), rabbit anti-HA monoclonal antibody (ROCKLAND), and rabbit anti-α-tubulin antibody (MBL). The corresponding secondary antibodies used were HRP-conjugated goat anti-mouse IgG (abcam), HRP-conjugated goat anti-rabbit IgG (abcam), and HRP-conjugated goat anti-chicken IgY (abcam).

### Minigenome reporter assay

MARV minigenome assays were performed as previously described (Wenigenrath *et al*., 2010). Briefly, plasmids for the minigenome assay (500 ng of pCAGGS-NP, 100 ng of pCAGGS-VP35, 100 ng of pCAGGS-VP30, and 1000 ng of pCAGGS-L, 500 ng of a MARV-specific minigenome encoding the *Renilla* luciferase reporter gene, and 500 ng of pCAGGS-T7 polymerase, and 50 ng of pGL-encoding firefly luciferase reporter gene for normalization) were transfected into HEK293 cells. EBOV minigenome assays were performed as previously described (Hoenen *et al*, 2010). Briefly, plasmids for minigenome assays (125 ng of pCAGGS-NP, 100 ng of pCAGGS-VP35, 100 ng of pCAGGS-VP30, 1000 ng of pCAGGS-L, with 250 ng of EBOV-specific minigenome encoding the *Renilla* luciferase reporter gene, 250 ng of pCAGGS-T7 polymerase, and 50 ng of pGL-encoding firefly luciferase reporter gene for normalization) were transfected into HEK293 cells. The cells were lysed and subjected to a luciferase reporter assay (PJK).

### Live cell imaging microscopy

A total 8 × 10^4^, 4 × 10^4^, or 2 × 10^4^ Huh-7 cells were seeded onto a µ-Dish 35 mm, µ-Slide 4 well, or 8 wells (ibidi) and incubated in DMEM/PS/Q with 10% FBS. To observe MARV nucleocapsid transport, cells were infected with a multiplicity of infection (MOI) of 1. The inoculum was replaced with fresh medium at 1 h post-infection (Dolnik *et al*., 2014). Subsequently, 500 ng of DNA encoding green fluorescent fusion protein was transfected. To observe MARV NCLS transport, each well was transfected with the following plasmids encoding MARV proteins: (500 ng of pCAGGS-NP, 100 ng of pCAGGS-VP35, 100 ng of pCAGGS-VP30^GFP^, and 100 ng of pCAGGS-VP24), together with MARV minigenome-expressing plasmid and T7 polymerase-coding plasmid (Hoenen *et al*., 2011; Wenigenrath *et al*., 2010). To observe EBOV NCLS transport, each well was transfected with the following plasmids encoding EBOV proteins: (500 ng of pCAGGS-NP, 200 ng of pCAGGS-VP35, 200 ng of pCAGGS-VP24, and 200 ng of pCAGGS-VP30-GFP). The inoculum was removed at 1 h p.t., and CO_2_-independent Leibovitz’s medium (Life Technologies) with PS/Q, non-essential amino acid solution, and 3–20 % (v/v) FBS was added. Live-cell time-lapse experiments were recorded with a Nikon ECLIPSE TE2000-E using a 63 × oil objective, GE Healthcare Delta Vision Elite using a 60× oil objective, Keyence BZ-X810 microscope using a 100× oil objective under BSL-2, and Leica DMI6000B under BSL-4 using a 63× oil objective equipped with a remote-control device to operate the microscope outside the BSL-4 facility (Schudt *et al*., 2013).

### Treatment of cells with cytoskeleton-modulating drugs

Cells were treated with 15 μM nocodazole (Sigma), 0.3 μM cytochalasin D (Sigma), or 0.15% dimethyl sulfoxide (DMSO; Sigma), as previously described (Schudt *et al*., 2013). The chemicals were added to the cell culture medium 3 h prior to observation.

### Image processing and analysis

The acquired pictures and movie sequences were processed using the Imaris tracking module (Bitplane; Oxford Instruments, Abingdon, UK) (Grikscheit *et al*, 2020). The size of spots > 1 μm for nucleocapsids and 0.5 μm for NCLSs were collected. Subsequently, “Quality” based optimization of the detected spots was performed. The tracking algorithm of “Autoregressive Motion” was applied for tracking with a “Maximum Distance” of 1 and “Maximum Gap” of 1 with a filling of gaps with detected objects. Detected trajectories of length > 5 μm, duration > 15 s, and track straightness > 0.2 were processed for the analyses. Moving signals were collected from approximately 10 cells derived from three independent experiments.

### Co-immunoprecipitation analysis

Co-immunoprecipitation assays were performed as previously described (Takamatsu *et al*., 2020a). For SDS-PAGE, elution was achieved with 70 μl of SDS sample buffer (Fujifilm) and subjected to gel electrophoresis and western blot analysis.

### Ultrathin section electron microscopy

Huh-7 cells were seeded on a 12-well plate, transfected with 1 μg NP or NP mutant-encoding plasmids, 0.5 μg VP24, and 0.5 μg VP35-encoding plasmids. At 48 h post-transfection, cells were fixed with aldehydes, postfixed with 1 % osmium tetroxide, dehydrated in a graded ethanol series, and embedded in EPON 812 (TAAB, Berks, UK). Ultrathin sections were stained with uranyl acetate and lead citrate and observed under a Hitachi HT-7700 microscope operated at 80 kV (Hitachi Hi-Tech, Tokyo, Japan) (Fujita-Fujiharu *et al*., 2025).

### VLP assay

Virus like particle (VLP) assays were conducted following established protocols (Biedenkopf & Hoenen, 2017; Hoenen *et al*., 2006; Wenigenrath *et al*., 2010), with minor modifications. HEK293 cells were seeded in six-well plates and transfected with plasmids encoding all structural proteins and reporter genes for either MARV (500 ng of pCAGGS-NP, 100 ng of VP35, 500 ng of VP40, 500 ng of GP, 100 ng of VP30, 100 ng of VP24, 1000 ng of L, 500 ng of pANDY-3M5M, 500 ng of pCAGGS-T7 polymerase) or EBOV (125 ng of pCAGGS-NP, 100 ng of VP35, 250 ng of VP40, 250 ng of GP, 100 ng of VP30, 100 ng of VP24, 1000 ng of L, 250 ng of pANDY-3E5E, 250 ng of pCAGGS-T7 polymerase). Plasmids encoding the firefly luciferase reporter gene were used for normalization. After 72 hours post-transfection, culture supernatants were collected, and VLPs were purified by ultracentrifugation using a 20% sucrose cushion. VLPs were analyzed for VP30 incorporation using proteinase K digestion assay (Wenigenrath *et al*., 2010).

### Alphafold-Multimer prediction

To predict the complex structure of NP-PPxPxY peptide and VP35 or VP24 in MARV or EBOV, we employed AlphaFold structural predictions. The amino acid sequences of EBOV and MARV proteins were obtained from GenBank (ID: EBOV: NC_002549, MARV: DQ217792.2). The structural complex of EBOV and MARV NPs with each peptide sequences were predicted in AlphaFold2 and AlphaFold3 software (Abramson *et al*., 2024; Jumper *et al*., 2021). The most confidently predicted structures (high pLDDT values) were visualized and assessed using UCSF Chimera software (Pettersen *et al*, 2004).

### Statistical analysis

Data represent the mean values and standard deviations from at least three independent experiments. Statistical analyses were performed using GraphPad Prism (version 8.0). Normally distributed samples were analyzed using the Student’s *t*-test. Statistically significant differences are indicated by asterisks (*, *P* < 0.05; **, *P* < 0.01; ***, *P* < 0.001; ****, *P* < 0.0001).

## Acknowledgments

The authors are grateful to Dr. Stephan Becker of Philipps-Universität Marburg, Germany; Dr. Hideki Ebihara; Dr. Masayuki Saijo; Dr. Masayuki Shimojima; Dr. Tomoki Yoshikawa; and Dr. Takeshi Kurosu for fruitful discussions. The authors also thank Susanne Berghoefer, Dirk Becker, Astrid Herwig, and Katharina Kowalski of Philipps-Universität Marburg, Germany; and Momoko Ogata, Satoko Sugimoto, and Masayasu Misu of the National Institute of Infectious Diseases, Japan; as well as all members of the Department of Virology at the Institute of Tropical Medicine, Nagasaki University, Japan, for their technical assistance. BSL-4 work would not have been possible without the supervision of Markus Eickmann and the technical support of Michael Schmidt and Gotthard Ludwig. This study was supported by Japan Society for the Promotion of Science, JSPS Grant numbers 18J01631, 19K16666, and 21K07059, 22KK0115; AMED under Grant Number JP23fm0208101, JP22fm0208101, JP23fk0108656, JP23wm0125006, JP22wm0325023, JP21wm0325023, and JP20wm0323023; Ichiro Kanehara Foundation for the Promotion of Medical Science & Medical Care; MSD Life Science Foundation ID-022; Japan Research Foundation for Clinical Pharmacology; TERUMO LIFE SCIENCE FOUNDATION; Waksman Foundation research grant; Novartis foundation research grant; Naito foundation grant; Takeda Science Foundation; Kurozumi Medical Foundation; Asahi Glass Foundation; Joint Usage/Research Center on Tropical Disease (2019-Ippan-24, 2020-Ippan-28, and 2021-Ippan-21); Joint Usage/Research Center program of Institute for Frontier Life and Medical Sciences Kyoto University (2020, 2021) (to Y.T.); AMED, Research Program on Emerging and Re-emerging Infectious Diseases (24fm0208101j0008, JP23fm0208101 and 22fk0108552h0001); JSPS Core-to-Core Program A, Advanced Research Networks (JPJSCCA20190008); JSPS Grant-in-Aid for Challenging Exploratory Research (22K19431); Grant for Joint Research Project of the Institute of Medical Science, University of Tokyo; Joint Usage/Research Center program of Institute for Frontier Life and Medical Sciences Kyoto University; Joint Usage/Research Center on Tropical Disease, Nagasaki University; and the Takeda Science Foundation (to T.N.).

## Materials availability

All plasmids, resources, and reagents generated in this study are available upon request and will be fulfilled by the lead contact, Yuki Takamatsu

## Data availability

Source images of IFA, Live-cell imaging, and TEM experiments were deposited in the BioImage Archive under accession number XXXX.

## Author contributions

**Yuki Takamatsu**: Conceptualization; Resources; Data curation; Formal analysis; Supervision; Funding acquisition; Visualization; Writing—original draft; Writing—review and editing; designed the experiments; performed and analysis for IFA, Live-cell imaging, TEM, IP, VLP, BSL-4 works. **Olga Dolnik**: Data analyses; Supporting BSL-4 works; Writing—review and editing; Supervision. **Ai Hirabayashi**: Performing TEM; Writing—review and editing. **Kenta Okamoto**: Prediction of structures; Resources; Data analysis; Writing—review and editing. **Tomomi Kurashige**: Preparation and analysis for molecular cloning and reporter assays; Writing—review and editing. **Hu Shangfan**: Preparation and analysis for molecular cloning; Writing—review and editing. **Catarina Oda Harumi:** Preparation and analysis for IFA and IP; Writing—review and editing. **Takeshi Noda**: Resources; Data curation; formal analysis; Supervision; Funding acquisition; writing—reviewing and editing.

